# Single-molecule assay reveals the impact of composition, RNA duplex, and inhibitors on the binding dynamics of SARS-CoV-2 polymerase complex

**DOI:** 10.1101/2025.01.10.632212

**Authors:** Terri C. Lovell, Heidi A. F. Dewling, Cynthia X. Li, Hery W. Lee, Calvin J. Gordon, Dana Kocincova, Maulik D. Badmalia, Egor P. Tchesnokov, Matthias Götte, Gonzalo Cosa

**Affiliations:** Department of Chemistry and Quebec Center for Advanced Materials (QCAM), McGill University, 801 Sherbrooke Street West, Montreal, QC, H3A 0B8, Canada; Department of Medical Microbiology and Immunology, University of Alberta, Edmonton, AB, T6G 2E1, Canada

**Keywords:** nucleoside inhibitor, non-nucleoside inhibitor, RNA dependent RNA polymerase, imaging, protein induced fluorescence enhancement

## Abstract

The genome replication of SARS-CoV-2, the causative agent of COVID-19, involves a multi-subunit replication complex consisting of non-structural proteins (nsps) 12, 7 and 8. While the structure of this complex is known, the dynamic behavior of the subunits interacting with RNA is missing. Here we report a single-molecule protein-induced fluorescence enhancement (SM-PIFE) assay to monitor binding dynamics between the reconstituted or co-expressed replication complex and RNA. Increasing binding times were observed, in this order, with nsp7 (none) nsp8 and nsp12, in nsp8-nsp12 mixtures and in reconstituted mixtures bearing all three proteins. Unstable, transient, and stable binding modes were recorded in the latter case, indicating that complexation is dynamic, and the correct conformation must be achieved before stable RNA binding can occur. Notably, the co-expressed protein yields mostly stable binding even at low concentrations, while the reconstituted proteins exhibit unstable binding indicating inefficient complexation with reduced protein. The SM-PIFE assay distinguishes inhibitors that impact protein binding from those that prevent replication, as demonstrated with suramin and remdesivir, respectively. The data reveals a correlation between binding lifetime/affinity, and protein activity, and underscores differences between co-expressed vs reconstituted mixtures, suggesting the existence of trapped conformations that may not evolve to productive binding.

## Introduction

In March 2020, the World Health Organization declared the novel coronavirus (COVID-19) outbreak as a global pandemic. Since then, over 7 million COVID-19 related deaths have been reported. COVID-19 is caused by SARS-CoV-2, a virus belonging to the *Coronaviridae* family. This is a positive-strand RNA virus containing a ∼30 kilobase single stranded viral RNA genome, making it one of the most substantial viral genomes.(1) Viral entry is mediated by spike proteins through receptor-mediated endocytosis.(2, 3) The viral genomic RNA is released, and the host cell machinery translates the RNA into two viral polyproteins (pp1a and pp1ab) required for viral replication and transcription. pp1a is processed into non-structural proteins nsp1-nsp11 and pp1ab is processed into nsp1-10 and nsp12-16.(4) Precursor polyproteins are cleaved by two proteases; the papain-like proteases (PLpro), encoded within nsp3, and a chymotrypsin-like protease, the main protease (Mpro), encoded by nsp5.(5)

The main nsps involved in viral genome replication and transcription of SARS-CoV-2 are nsp7, nsp8 and nsp12. Notably, nsp12 is an RNA-dependent RNA polymerase (RdRp), which catalyzes RNA replication.(6, 7). RdRp copies the (+)-strand RNA genome into the (-)-strand, which then serves as a template for (+)-strand RNA production.(8, 9). Co-factors nsp7 and nsp8,(10) are required for processive RNA synthesis activity.(1) Numerous structural and bulk studies on the RdRp nsp12 and its cofactors nsp7 and nsp8 have been conducted, including in the presence of inhibitors such as remdesivir. (6, 7, 10-20) Subissi et al. first determined the requirement of nsp7 and nsp8 for replication in SARS-CoV, which is highly similar in sequence and structure to SARS-CoV-2.(18) The structure of nsp12 bound to nsp7 and nsp8 was resolved by Kirchdoerfer and Ward using single particle cryoEM. (7) This revealed the conserved structure of the RdRp when compared to other viral polymerases. The core complex while bound to RNA was then resolved, illustrating how the RNA sits in a “right hand” RdRp domain (residues S367-F920), composed of fingers (L366-A581 and K621-G679), palm (T582-P620 and T680-Q815) and thumb (H816-E920) subdomains. (6, 7, 11-16) This study also showed long helical extensions as nsp8 protrude along exiting RNA, that can account for the processivity of the RdRp. Furthermore, the active structure with an incorporated molecule of remdesivir was determined, providing valuable insight into its mechanism for polymerase stalling. Selectivity for remdesivir triphosphate over ATP, its natural counterpart, was determined through an analysis of the active site for NTP incorporation. (19) Suramin, a 100-year-old inhibitor, was also found to inhibit the SARS-CoV-2 RdRp by blocking the RNA binding site. (20) These structural studies provide foundational knowledge of nsp12 complexation with nsp7 and nsp8, how they are bound to RNA duplexes, and how this is impeded by inhibitors such as remdesivir and suramin. However, these results are a static representation of the RdRp complex, and do not reflect the dynamic nature of polymerase-RNA binding interactions.

Currently, a real-time analysis of the interaction dynamics and associated affinity of the viral replication complex with its RNA duplex is missing. Knowledge is required on ***i***. the ability of individual elements (nsp7, nsp8 and nsp12) to bind and their association and dissociation rates, ***ii***. the impact of nsp7, nsp8 and nsp12 altered stoichiometry on these rates and overall affinity, ***iii***. what impact the method of the protein expression (individually expressed vs co-expressed) plays, ***iv***. the influence of RNA secondary structure, and ***v***. inhibitor impacts on the binding affinity. This understanding is necessary for developing drug therapies, where real-time monitoring of protein-RNA dynamics is required to understand replication mechanisms and screen for potential inhibitors.

Herein we exploit a SM fluorescence (21-25) assay relying on protein induced fluorescence enhancement (SM-PIFE)(26), to study the binding dynamics of the key nsps involved in SARS-CoV-2 viral genome replication. Our work illustrates that while nsp7 lacks any binding to RNA duplexes alone, nsp8 shows weak affinity and nsp12 a rather tight binding, where a combination of the latter two results in a synergistic activity with rapid and stable binding to RNA. PIFE binding assays further show that increasing ratios of accessory proteins nsp7 and nsp8 to nsp 12 result in enhanced binding and correlates with increased RdRp activity. In turn, co-expressed nsp7/8/12 shows optimal binding even at low concentrations, in contrast to separately expressed nsp7, nsp8, and nsp12 reconstituted at time of experiment. The SM-PIFE assay can further decipher between non-nucleoside inhibitors (NNIs) and nucleoside analog polymerase inhibitors (NAPIs) and their impact on assembled replication duplexes. Together, the single molecule data shows that different subunits have differing binding affinities and that either the co-expressed complex or the complex formed by nsp12 with an excess of 8 and 7 show maximum binding, and optimal primer extension at 100% and 35%, respectively. The SM-PIFE assay reported herein further permits readily monitoring the impact that order of addition of inhibitors have on the complex binding dynamics.

## Materials and Methods

### Slide preparation and surface immobilization

Acetone (HPLC grade), Premium Cover Glass slides (25×25 mm), sulfuric acid, and sodium hydroxide were purchased from Fisher. Dimethyl sulfoxide (anhydrous, ≥99.9), Wheaton Coplin Staining Jars were purchased from SigmaAldrich. Hydrogen peroxide (30% v/v) was purchased from ACP Chemicals. Silicon molds, polycarbonate imaging chambers (Hybriwell^TM^), and ports (press fit tubing connectors) were purchased from Grace Bio-Labs, Inc. Double sided tape to affix ports was purchased from Scor-pal. Poly(ethylene glycol) polymers mPEG-Silane (MW 5000) and Biotin-PEG-Silane (MW 5000) were purchased from Laysan Bio Inc. Polymers were dissolved in anhydrous dimethyl sulfoxide (DMSO) and a 25% (w/w) PEG-Silane and 0.25% (w/w) Bio-PEG-Sil solution was made. Glass coverslips were cleaned, etched, and passivated as previously described. (27) Polycarbonate imaging chambers were installed with a silicone tubing connector on either end to allow solutions to be flowed across the slide surface. The surface was incubated with a 0.2 mg/mL streptavidin solution (prepared in 50 mM HEPES) for 5 minutes. Unbound streptavidin was rinsed away with 50mM HEPES. Following streptavidin incubation, a 25 pM RNA solution (in 50 mM HEPES buffer pH 8.0) was flowed through the chamber and oligomers were immobilized to the PEG-coated glass coverslips via biotin-streptavidin interactions. Unbound RNA was gently rinsed with additional 50 mM HEPES. In experiments with inhibitors, a 100 nM biotin solution was added to prevent unwanted interactions with the slide surface.

### Nucleic acids

RNA template and primer were purchased from Integrated DNA Technologies. Template and primer strands were annealed by incubating at 95°C for 2 min followed by a gradual cooling (2°C/min) to 25°C. The RNA template with the longer 25 nt overhang is a Cy3-tagged 55mer (/5Cy3N/GCA-CUU-AGA-UAU-GAC-UCG-UUC-UGC-AGG-CCA-GUU-AAU-AAC-GUC-UAA-GAC-ACA-GAU-C/3) and the primer strand is a biotinylated 30mer (/5Bio/GAU-CUG-UGU-CUU-AGA-CGU-UAU-UAA-CUG-GCC/3). The RNA duplex with the shorter 20 nt overhang (duplex 2, below) consisted of template strand Cy3-tagged 58mer (/5Cy3N/CCA-CAC-AAC-ACC-UAC-GGC-AAU-GGA-GCG-CUG-GCA-GCG-GUU-AAC-GUC-UAA-GAC-ACA-GAU-C/3) and primer strand biotinylated 38mer (/5Bio/GAU-CUG-UGU-CUU-AGA-CGU-UAA-CCG-CUG-CCA-GCG-CUC-CA/3).

### Proteins

Expression and purification of SARS-CoV-2 RNA-dependent RNA polymerase complex consisting of nsp7, nsp8, nsp12 (RdRp) have been described previously (28). Briefly, the pFastBac-1 (Invitrogen, Burlington, Ontario, Canada) plasmid with the codon-optimized synthetic DNA sequences (GenScript, Piscataway, NJ) coding for a portion of 1ab polyprotein of SARS-CoV-2 (NCBI: QHD43415.1) containing only nsp5, nsp7, nsp8, and nsp12 were used as a starting material for protein expression in insect cells (Sf9, Invitrogen). We employed the MultiBac (Geneva Biotech, Indianapolis, IN) system for protein expression in insect cells (Sf9, Invitrogen) according to published protocols (46, 47). SARS-CoV-2 protein complexes were purified using nickel-nitrilotriacetic acid affinity chromatography of the nsp8 N-terminal eight-histidine tag according to the manufacturer’s specifications (Thermo Scientific).The individual components of SARS-CoV-2 RdRp were expressed as follows: nsp12 with a C-terminal eight-histidine tag was expressed in insect cells by following the procedure outlined above; nsp7 and nsp8 with the N- and C-terminal eight-histidine tags, respectively, were expressed in *E*.*coli* from pET-15b plasmids containing the respective genes that were codon-optimized for expression in *E*.*coli* (GenScript, Piscataway, NJ). All three proteins were individually purified using the eight-histidine tags according to the manufacturer’s specifications (Thermo Scientific). RdRp complex was reconstituted by mixing the individual components at indicated proportions with respect to final nsp12 concentration after mixing all the components.

### Gel-based assay for RNA synthesis by SARS-CoV-2 RdRp complex

RNA primer/template sequences used to monitor the RNA synthesis are shown in Figure 3A. A 38-nt primer (biotinylated at the 5’-end) was heat-annealed to a 58-nt template (Cy-3 labeled at the 5’-end) in 25 mM Tris-HCl (pH8) buffer supplemented with 50 mM NaCl. RNA synthesis assay consisted of pre-incubating 150 nM of SARS-CoV-2 RdRp complex with 25 nM primer/template in the presence of 0.1 μM [α-^32^P]-UTP, 0.1 μM (or 1 μM for order of addition experiments) NTP mix (ATP, CTP, GTP) and 0.2 mM EDTA in 25 mM Tris-HCl (pH 8) for 10 minutes at 30 °C. The RNA synthesis was initiated by addition of 5 mM MgCl_2_. Reactions (15 μL) were incubated for 90 seconds at 30 °C and then stopped by the addition of 45 μL of formamide/EDTA (25 mM) mixture and incubated at 95 °C for 10 min. 3 μL reaction samples were subjected to denaturing 8 M urea 20% polyacrylamide gel electrophoresis to resolve products of RNA synthesis followed by signal quantification (ImageQuant 5.2, GE Healthcare Bio-Sciences, Uppsala, Sweden) through phosphorimaging (Amerhsham Typhoon 5, Cytivia, Marlborough, MA, USA).

### Buffers and reagents

MgCl_2_ 1.0 M, HEPES 1.0 M buffer solution pH 8.0, and Molecular Biology Grade HyClone were acquired from ThermoFisher Scientific (South Logan, UT). Unlabeled streptavidin (S888) and nucleoside triphosphate (CTP, ATP, GTP and UTP) 100 mM solutions were purchased from Thermofisher Scientific (Invitrogen). D-(+)-glucose, glucose oxidase (from Aspergillus niger, type VII) and biotin (HPLC grade lyophilized powder) were purchased from Sigma-Aldrich. Suramin was purchased from EMD Millipore. Remdesivir (active triphosphate metabolite GS-443902 trisodium) was purchased from MedChemExpress. Glycerol monothioglycolate (GMTG), 80%, was purchased from Evans Chemetics.

### Anti-fading solution

Experiments were carried out in the presence of an oxygen scavenger solution consisting of D-(+)-glucose (3% w/v) and glucose oxidase (165 units/mL), as well as 21 mM triplet state quencher GMTG (Glycerol Monothioglycolate 80%) and 5 mM MgCl_2_ in 50 mM pH 8.0 HEPES. The first image was acquired 10 min after introducing the oxygen removal system. Experiments were conducted at room temperature (23 °C).

### TIRF microscopy

Fluorescence imaging was carried out using an inverted Nikon Eclipse Ti microscope equipped with the perfect focus system (PFS) implementing an objective-type TIRF configuration with a Nikon TIRF illuminator and an oil-immersion objective (CFI SR Apo TIRF 100×, numerical aperture (NA) 1.49). With these settings, the 561nm (149 W/cm2) laser was used for excitation (Agilent MLC400B Monolithic Laser Combiner): 561 nm, based on a power measured at the objective and a beam sized as the FOV ∼ 82 x 82 μm ). The laser beam was passed through a multiband cleanup filter (ZET405/488/561/647x, Chroma Technology) and coupled into the microscope objective using a multiband beam splitter ZT405/488/561/647rpc (Chroma Technology). Fluorescence light was spectrally filtered with an emission filter ZET405/488/561/647m (Chroma Technology). For Cy3 imaging, an additional emission filter ET600/50m (Chroma Technology) was used. All movies were recorded onto a 512 x 512 pixel region of a back-illuminated electron-multiplying charge-coupled device (EMCCD) camera (iXon X3 DU-897-CS0-#BV, Andor Technology), with a pixel size of 16 μm. Given the 100× magnification used, the effective pixel size was 160 nm. The camera was controlled using Micro-Manager Software (Micro-Manager 1.4.13, San Francisco, CA, USA), capturing 16-bit 512 x 512 pixel images with an exposure time of 50 ms. The microscope and laser powers were controlled using the software NIS element from Nikon.

### Statistical analysis

Experiments were performed either in duplicate or triplicate, where the results of each trial are listed individually in the Supporting Information. The sample size of each trial is denoted by n. Error bars represent means ± standard deviation.

## Results and Discussion

### SM-PIFE assay to elucidate viral protein interactions with RNA duplexes

To evaluate protein binding to RNA, a single-molecule based protein induced fluorescence enhancement (SM-PIFE) assay was designed (**Figure 1A**). In this assay, protein binding to a Cy3-tagged RNA leads to an emission enhancement that is related to protein proximity. The enhancement is associated with a decrease in the non-radiative decay pathway of the fluorophore by preventing rotational deactivation (internal conversion) in the photoexcited fluorophore. A primer-template RNA duplex was designed where the primer strand (38-nt) was biotinylated at its 5’-end for surface immobilization. The template strand (58-nt or 63-nt long) had a 20 or 25 base pair (bp) single-stranded (ss) overhang bearing Cy3 at the 5′ end (**Figure 1A**). The dsRNA region was designed long enough to accommodate binding of the fully assembled nsp7/8/12 complex. Following annealing, the primer-template RNA duplex was immobilized to a polyethylene glycol (PEG) passivated glass coverslip using biotin-streptavidin interactions.(27) Intensity-time trajectories were recorded utilizing a total internal reflection fluorescence (TIRF) microscope equipped with a 561 nm, 10 mW output continuous wave laser for excitation and an EMCCD camera for image capture. To ensure rapid data collection while preserving signal sensitivity, imaging was conducted at a rate of 20 frames/second. Experiments were performed in HEPES buffer (50 mM, pH 8), 5 mM Mg^2+^, under oxygen-depleted conditions (to maximize fluorophore stability) using the glucose/glucose oxidase system (GlOx). While catalase is typically utilized in the GlOx deoxygenation system, it was deliberately excluded in this study due to the presence of RNases. Glycerol monothioglycolate was used as a triplet state quencher (TSQ). Two separate enzyme preparations, which differed in their production strategy, were utilized. One method involved insect cell (*Sf9*) co-expression of the nsp7, nsp8, and nsp12 proteins as part of the polyprotein and in frame with nsp5 protease located upstream of the polyprotein (28). Thus, the expressed polyprotein is post-translationally cleaved into the individual components and the formed nsp(7:8:12) complex is purified using Ni-NTA affinity chromatography on the nsp8 N-terminal histidine tag. Alternatively, the individual components were expressed separately in *E*.*coli* (nsp7, nsp8) and *Sf9* (nsp12), purified and reconstituted according to the indicated stoichiometry of the individual components.

**Figure 1.**
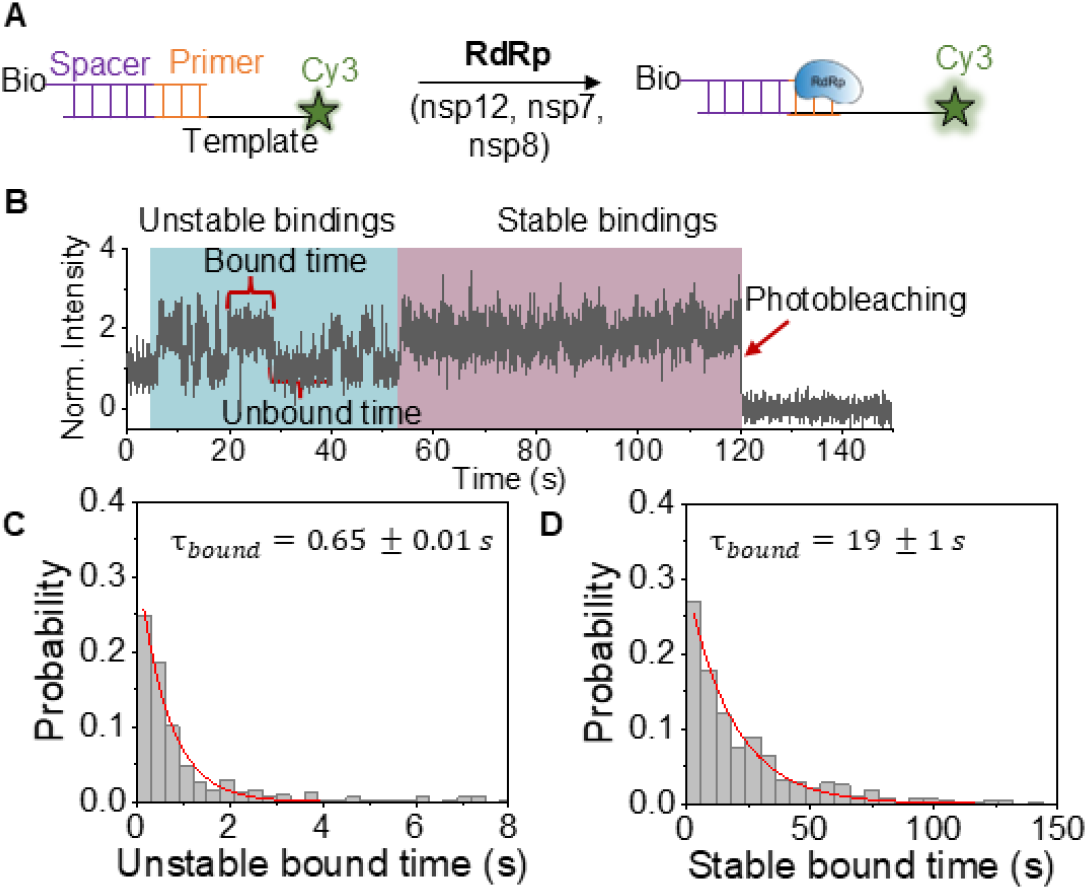
(**A**) Protein Induced Fluorescence Enhancement (PIFE) assay used to monitor protein binding to RNA. (**B**) Representative single-molecule trajectory showing protein binding, time bound to RNA (τ_bound_, unbound time (τ_unbound_), and photobleaching time for separately expressed nsp7, nsp8, nsp12. (**C**) Probability vs on time histogram for all molecules to determine τ_bound_ for unstable and (**D**) stable binding events.

After recording the initial 12 seconds (240 frames) in a GlOx/TSQ solution without nsps, a solution containing one or more nsp7, nsp8, and/or nsp12, or the co-expressed protein was manually injected. Subsequent images were captured over time until most fluorophores exhibited photobleaching, evident in SM-trajectories as a single drop in intensity to background levels. The stochastic association and dissociation of proteins resulted in changes to the intensity trajectory over time. Protein binding to the single RNA duplexes resulted in a ∼2-fold enhancement in Cy3 fluorescence (**Figure 1B**). Protein dissociation in turn resulted in fluorescence decrease to the initial (pre-nsp addition) Cy3 intensity (**Figure 1B**). From the intensity-time trajectories, the protein bound (on) and unbound (off) dwell times were extracted for every single molecule analyzed in a given condition (protein concentration, inhibitors, etc). The unbound and bound times from all molecules analyzed in a given condition were combined to create a probability distribution histogram (**Figures 1C and 1D**). Fitting the histograms to a single exponential allowed lifetime determination for protein association (τ_unbound_, obtained from the distribution of the “off” dwell times), and protein dissociation (τ_bound_, obtained from the distribution of the “on” dwell times). The photobleaching time, effectively the observation time amenable to our experiments, was also extracted from the distribution of times recorded under each experimental condition. Importantly, RNA duplexes that in the presence of protein mixtures interconverted stochastically between the two states (high - PIFE - and low intensity) are herein described as exhibiting “unstable” protein binding. When unstable binding was followed by stable binding, these trajectories were referred to as “transient” binding. The lack of PIFE in turn signals no protein binding. Duplexes that exhibited a single binding event lasting until photobleaching precluded further investigation (only shown when nsp12 is present, *vide infra*), are referred to as showing “stable” protein binding. These stable binding events were confirmed via an experiment where immobilized RNA duplexes were monitored initially and following 30 min incubation (in the dark to prevent photobleaching) with protein. Here a total of 5 s were recorded following protein addition (100 frames) to position and analyze the initial protein binding response. Following the 30 min incubation, duplexes were imaged again. Duplexes that initially showed bound proteins remained bound to proteins, while duplexes lacking protein appeared bound to proteins after the incubation period (**Table S1**). Bindings that persisted until photobleaching, in the large majority of cases, were 100 times larger than the average unstable binding time. Therefore, events are classified as stable only if there is no unbinding event and their duration is 10 times longer than the average unstable binding time.

### nsp7, nsp8, and nsp12 reveal exhibit variable binding dynamics

We initially focused our attention on the binding kinetics and affinities of the individual components of the RdRp complex nsp7, nsp8, and nsp12. These proteins were expressed separately either in an *E. coli* (nsp7 and nsp8) or insect cells *Sf9* (nsp12) expression system.(28) SM analysis was conducted to understand how each nsp component individually contributes to the binding of the SARS-CoV-2 complex (**Figure 2**). By fitting probability histograms of both bound and unbound times, binding and equilibrium constants were calculated. The association rate constant (*k*_*a*_) was calculated using the average unbound time (τ_*unbound*_) and the given nsp protein concentration, as seen in Equation 1.

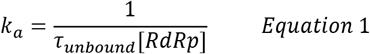

**Figure 2.**
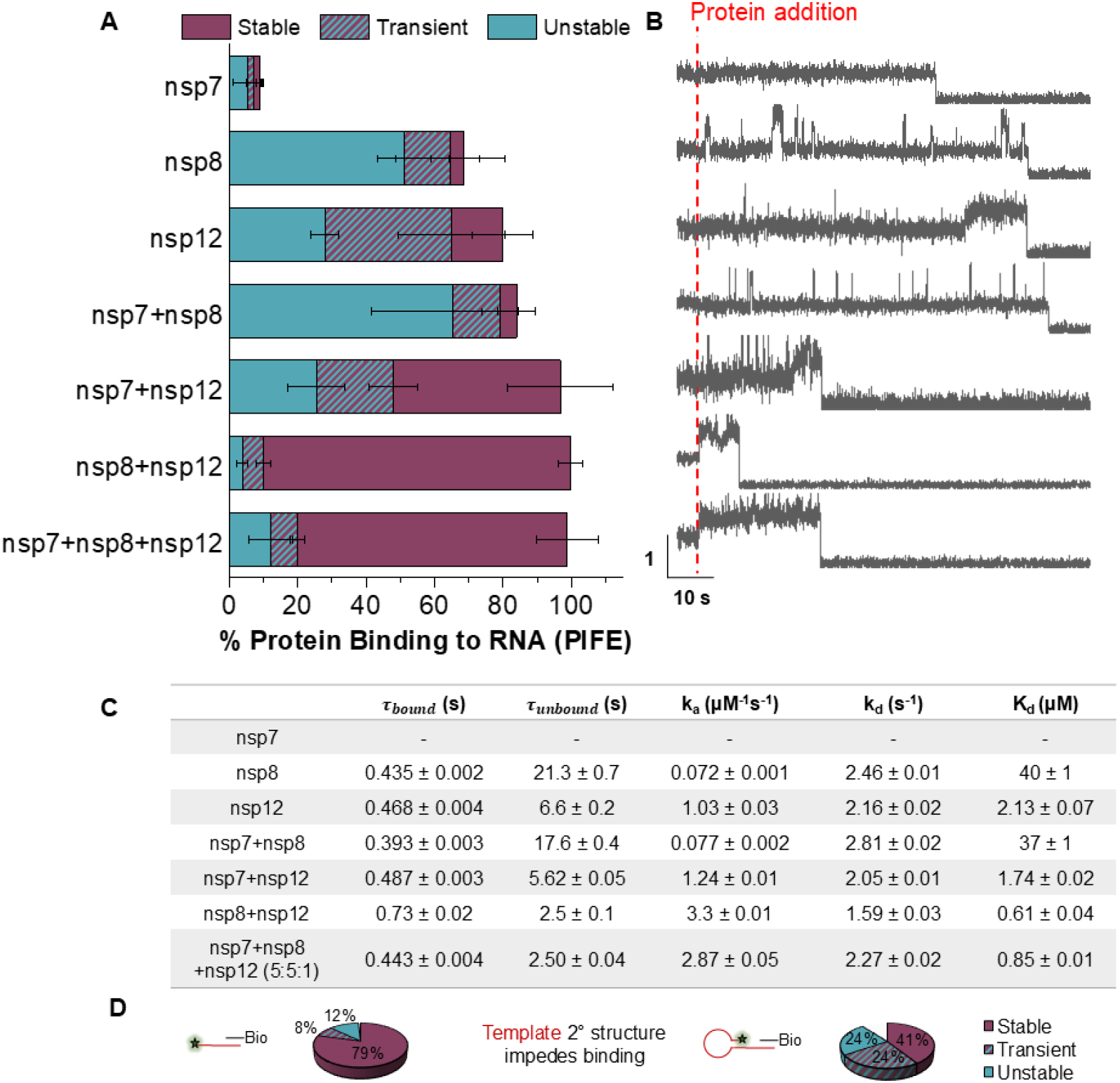
(**A**) The percent of molecules that showed protein binding to RNA (PIFE) broken into unstable, transient, and stable bindings for each RdRp component. (**B**) Corresponding representative SM trajectory of each RdRp component. (**C**) Average on time (τ_bound_), average off time (τ_unbound_), rate of association (k_a_), rate of dissociation (k_d_), and dissociation constant (K_d_) of different SARS-CoV-2 RdRp components. These values were calculated using only unstable bindings. (**D**) Cartoon representation of RNA with 2° hairpin structure and pie chart showing how the RNA structure impacts binding of reconstituted protein.

**Figure 3.**
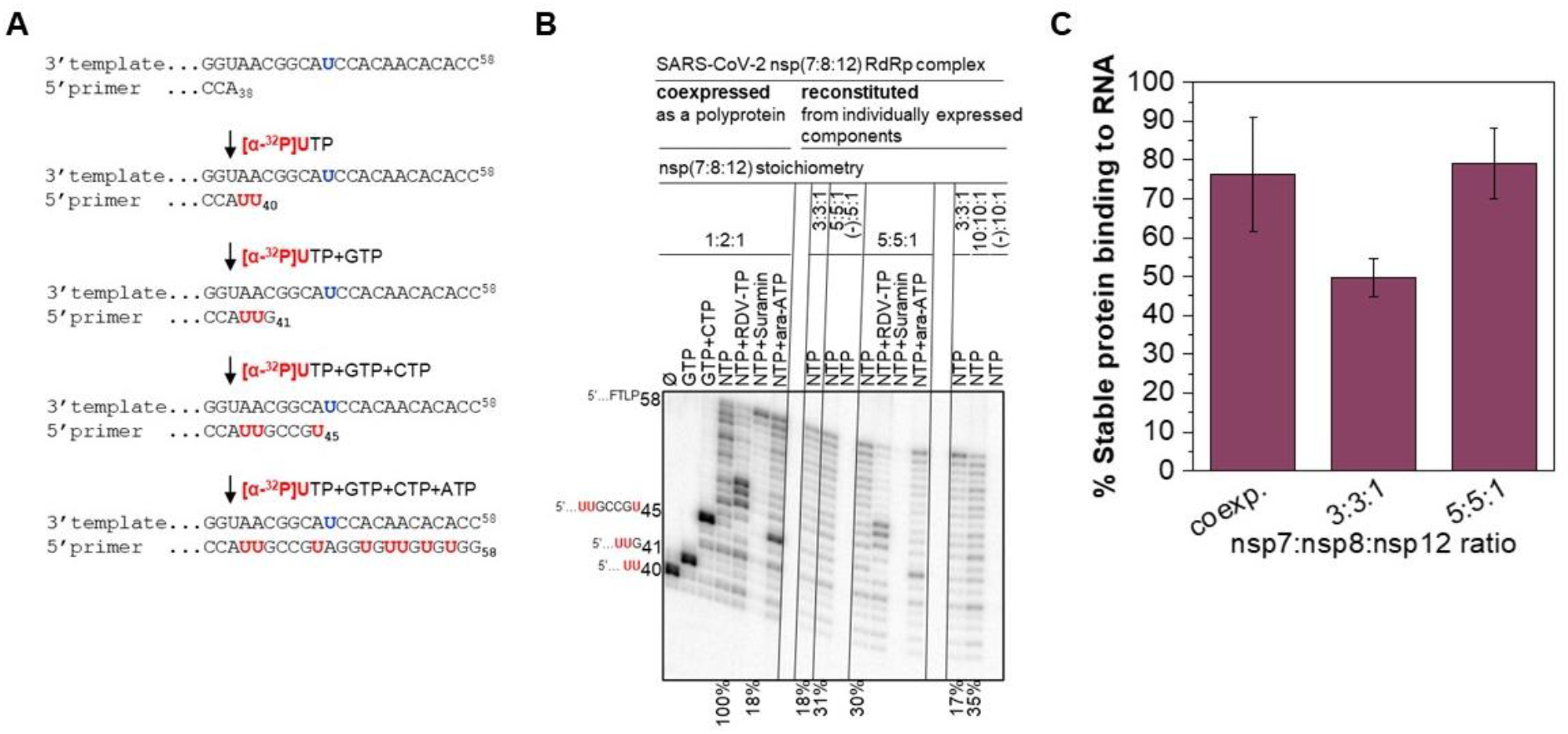
Comparison of RNA synthesis activity catalyzed by SARS-CoV-2 RdRp complexes co-expressed or reconstituted. Schematic representation of the RNA synthesis assay. For clarity only a segment of the sequence of the heat-annealed RNA primer/template system used to monitor the RNA synthesis is shown. Numbers in superscript format indicate the nucleotide length of the template. Numbers in subscript format indicate the nucleotide length of the primer and the expected products. Downward arrows illustrate the addition of specific NTP mixes to the reaction mixtures, which allows the formation of primer extension products with defined length. *Bold red*, illustrates a radiolabeled UTP added to the reaction mixture, or incorporated radiolabeled UMP during primer extension, which effectively labels the reaction products and allows signal detection and quantification. *Bold blue*, illustrates the template position opposite which remdesivir may be incorporated. (**B**) Denaturing PAGE migration pattern of the products of RNA synthesis along the primer/template shown in panel A. Ø, indicates that only radiolabeled [α-^32^P]UTP was added to the reaction mixture. GTP, or GTP+CTP indicate that the radiolabeled [α-^32^P]UTP was supplemented with the corresponding nucleoside triphosphates. NTP, a mixture of ATP, CTP, and GTP. RDV-TP, tri-phosphorylated remdesivir. FTLP, full template-length product. Sequences of letters shown to the left of the gel illustrate the nucleotide-extended primers in the presence of specific NTP combinations added to the reaction mixtures as per panel A. Numbers associated with the letter sequences illustrate the nucleotide size of the extended primers. Percent numbers shown below the gel indicate the relative values of the quantified total signal in the corresponding lane, a measure of the extent of the RNA synthesis. Difference in the values corresponding to the same reaction conditions illustrate the quantification error associated with sample loading. 100%, illustrates the lane (i.e. the reaction condition) that was used as a reference for comparisons. (**C**) Percent of stable binding of varying ratios of separately expressed SARS-CoV-2 RdRp components as determined from SM-PIFE experiments.

Similarly, the dissociation constant (*k*_*d*_) was calculated using the average bound time (τ_*bound*_), as seen in Equation 2.

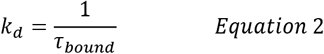

The equilibrium dissociation constant (*K*_*d*_) for the dissociation of protein from the RNA duplex was in turn established from the ratio of the dissociation and association rate constants according to Equation 3.

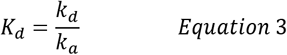

Among the individual components of the RdRp complex, nsp7 was rarely observed to bind (**Figure 2A**), which manifested as no change in fluorescence intensity along the single intensity-time trajectories recorded (**Figure 2B**). In very few trajectories, one sole unstable binding event was observed, likely an impurity in solution. The lack of PIFE signals indicates nsp7 does not bind to RNA on its own long enough to be recorded in a single frame (50 ms duration). In turn, for nsp8, the intensity-time trajectories showed several binding events, each extending for a few frames at a time, throughout the experiment. Specifically, 69% of the molecules showed unstable binding. Analyzing trajectory dwell times for these nsp8 unstable bindings revealed stochastic association and dissociation events, providing *k*_a_ = 0.072 ± 0.001 µM^-1^s^-1^ and K_d_ = 40 ± 1 µM. In 4% of the trajectories, the unstable events were succeeded by a final stable binding. Moving next to nsp12, while the intensity-time trajectories initially showed some unstable binding events, most molecules ultimately reached a stable binding configuration within a few seconds and before fluorophore photobleaching. The nsp12 exhibited the highest overall binding (86%), with only 26% of it being unstable, significantly less than in the case of nsp8 alone. The association rate constant and equilibrium dissociation constant for the unstable events were found to be *k*_a_ = 1.03 ± 0.03 µM^-1^s^-1^ and K_d_ = 2.13 ± 0.07 µ, respectively. These results indicate that both nsp12 and nsp8 contribute significantly to the binding of the complex with its RNA duplex.

We next compared mixtures containing any two of the three nsps involved in RdRp activity. Since nsp12 is the limiting factor for polymerization, its concentration was used when calculating *k*_*a*_ according to Equation 1. If nsp12 was not present, the concentration of nsp8 was used instead as it is the more critical accessory protein when it comes to binding, based on the above findings. In all cases the concentration of accessory components nsp7 and nsp8 was 5-fold greater than the concentration of nsp12.

The combination of nsp8 and nsp12 (5:1) exhibited an immediate intensity increase in the SM-trajectories, which persisted until photobleaching. This behavior was observed in 90% of SM-trajectories that exhibited PIFE. Moreover, the nsp8+nsp12 combination demonstrated the highest overall binding (PIFE) at 96%. For the unstable binding events, association and dissociation rate constants were found to be *k*_a_ = 3.3 ± 0.01 µM^-1^s^-1^ and k_d_ = 1.59 ± 0.03 s^-1^, which are the best for the three possible two-protein combinations tested. The trajectories of the nsp7+nsp8 experiments closely resembled those of nsp8 alone, where there are mostly unstable binding events. This is expected, as nsp7 does not bind. Furthermore, the association and dissociation rate constants of nsp7+nsp8 are within error of nsp8 (**Figure 2C**). A similar observation is made in the case of nsp7+nsp12, where SM-trajectories of nsp7+nsp12 show similarities to nsp12 alone. Here there are some unstable binding events as well as a significant portion of stable bindings. The association and dissociation rate constants for nsp7+nsp12 are similar to that of nsp12 alone, which is expected since nsp7 was found not to bind.

Subsequently, we tested a mixture with all three nsps present at a ratio of 5:5:1 nsp7:nsp8:nsp12. Upon protein addition there was an immediate increase in fluorescence intensity, reflecting rapid binding and affinity of the tetramer for the RNA duplex, which remained until photobleaching, and showed ∼79% stable binding. The overall percent PIFE and distribution of binding behavior was similar to nsp8+nsp12 (5:1). In turn, the presence of stable binding markedly decreased in the absence of either nsp12 or nsp8, to 5% and 49% in the nsp7+nsp8 (1:1) and nsp7+nsp12 (5:1) mixtures, respectively (**Figure 2**).Together, this data suggests that nsp8 and nsp12 play a significant role in the initial binding of the complex, while nsp12 is crucial for achieving stable binding and polymerization. Importantly, although nsp7 does not seem to have an impact on complex binding (nsp8+nsp12 versus nsp7+nsp8+nsp12 rate constants), it is required for polymerization (**Figure 3B**, (-):5:1 and (-):10:1).

Consistent with the nsp7, nsp8, nsp12 proposed binding mechanism, we also saw that the binding behavior of the protein is sensitive to RNA secondary structure. Thus, when an RNA duplex with a 25-nt overhang with self-complementarity, producing a secondary hairpin structure, was tested (**Figure 2D**), the number of molecules with stable binding went from 79% (RNA with no secondary structure) to 41% on the hairpin RNA. Furthermore, the number of transient events increased from 8% to 24% and there was an 11-fold increase in non-binding (1% to 11%). This indicates that the hairpin prevents the large complex from binding, where the high frequency of unstable binding events may be explained by the presence of one or multiple nsps binding to the RNA, rather than the entire nsp7-8-12 complex.

### Enhanced binding and RdRp activity observed with increasing ratios of nsp7 and nsp8 to nsp12

We next explored how varying the stoichiometry of individually expressed nsp7, nsp8, and nsp12 influences the association and dissociation rate constants and overall affinity of reconstituted mixtures (**Table S4**) and compared these results with the relative levels of the RNA synthesis catalyzed by the corresponding RdRp complexes in the gel-based biochemical assays (**Figure 3B**).

RNA synthesis catalyzed by either the co-expressed or the reconstituted RdRp complexes was monitored in the biochemical gel-based assays (**Figure 3B**), which rely on nucleotide incorporation into an RNA primer along the heat-annealed RNA template in the presence of magnesium ions as catalysts. Incorporation of radiolabeled [α-^32^P]-UTP allows the monitoring the formation of reaction products and their quantification through phosphorimaging. Thus, the quantified total signal in the lane may be used as a measure of the overall RNA synthesis activity. RdRp complexes co-expressed as a polyprotein incorporated the supplemented nucleotides into the RNA primer along the RNA template in a template base-specific manner (**Figure 3B**, first four lanes from the left). Supplementing the reaction mixtures with RDV-TP or ara-ATP resulted in signal accumulation after three nucleotide or right after the site of incorporation, respectively, which is consistent with the delayed-or immediate chain termination of primer extension. Suramin was used a positive control here and, as expected (20), inhibited RNA synthesis through competition with the RNA duplex as illustrated by the reduction in the overall signal in the lane by ∼80%. RdRp complexes that were reconstituted from the individual components according to the nsp(7:8:12) stoichiometry of 3:3:1, 5:5:1, and 10:10:1 exhibited similar patterns of RNA synthesis and its inhibition, albeit at the overall lower levels: ∼20, 30, and 35 % with respect to RNA synthesis by RdRp complex co-expressed as a polyprotein. The data also suggests that the maximal RNA synthesis activity of the reconstituted RdRp complex is obtained with 5:5:1 stoichiometry of the individual components and increasing the stoichiometry to 10:10:1 did not elevate the levels of RNA synthesis to the levels of RdRp complex of the co-expressed components. Omitting nsp7 during the reconstitution of the RdRp complex ablated RNA synthesis activity, thus illustrating its essential role for the formation of the catalytically active RdRp complex.

The enhanced RNA binding observed in these experiments correlated with the increase in the levels of RNA synthesis by the RdRp complexes reconstituted at the 3:3:1 and 5:5:1 stoichiometries. Altering the stoichiometric ratio of the RdRp components - from the 1:2:1 nsp7:nsp8:nsp12 found in the crystal structure of the core complex - was found to enhance protein binding to RNA (**Figure 3C**); where the highest affinity was found with the 3:3:1 or 5:5:1 stoichiometry of the components of the complex (k_a_ = 3.17 ± 0.05 µM^-1^s^-1^). Notably, the 5:5:1 stoichiometry exhibited a much higher percentage of stable bindings (79%) relative to the 3:3:1 ratio (50%), as shown in **Figure 3C**. This is consistent with a higher likelihood of formation of a stable complex with excess nsp7 and nsp8. The enhancement of RNA binding and the increase in RNA synthesis appeared to reach saturation in the presence of RdRp complexes of 5:5:1 stoichiometry.

### Co-expressed RdRp complex displays superior binding and RdRp activity

To establish whether the protein synthesis protocol impacts the affinity and activity of the replication complex, we next compared the binding of nsp7, nsp8, and nsp12 that were expressed separately and reconstituted *in vitro*, to co-expressed nsp7/8/12 using the insect cells Baculovirus expression system. (28) Here, we anticipated that formation of a productive complex would be favored in the co-expressed construct. We chose a 2:2:1 ratio of reconstituted nsp7, nsp8, and nsp12 because this ratio ensures there are enough accessory proteins for efficient complex formation, while preventing monomers from competing with tetramers.

Working with concentrations of 0.15 µM for reconstituted and co-expressed constructs, we observe that the amount of overall binding and stable binding of the co-expressed complex was similar to that of the reconstituted 2:2:1 complex (**Figure 4A**). To further investigate potential differences between individually expressed and co-expressed proteins at the SM-level, we reduced the concentration of proteins by 3-fold for both experiments with reconstituted and co-expressed constructs (**Figure 4B-C**). Here, we argued, the need to form a productive complex would negatively impact the binding of reconstituted mixtures, where a drop in association rate constants is expected. In turn, a drop in the concentration of co-expressed would not impact the structure or lead to a slowdown in the rate of binding. Consistent with our expectations, experiments under the new conditions showed that the association rate constant of co-expressed nsp7/8/12 remained unperturbed. In contrast, the association rate constant of the reconstituted nsps decreased 5-fold. Prior to dilution, both co-expressed and reconstituted proteins predominantly exhibited PIFE signals from stable binding, with the remaining 20% divided between transient and unstable bindings (**Figure 4A**). Dilution of the co-expressed nsp7/8/12 revealed that the majority of binding events remained stable or transient. Notably, dilution of the reconstituted proteins led to most binding events being unstable, with transient and stable bindings constituting the smallest portion. Likely these binding events originate from individual nsps or a combination of two nsps, indicating the formation of the replication complex is impaired under increased dilution.

**Figure 4.**
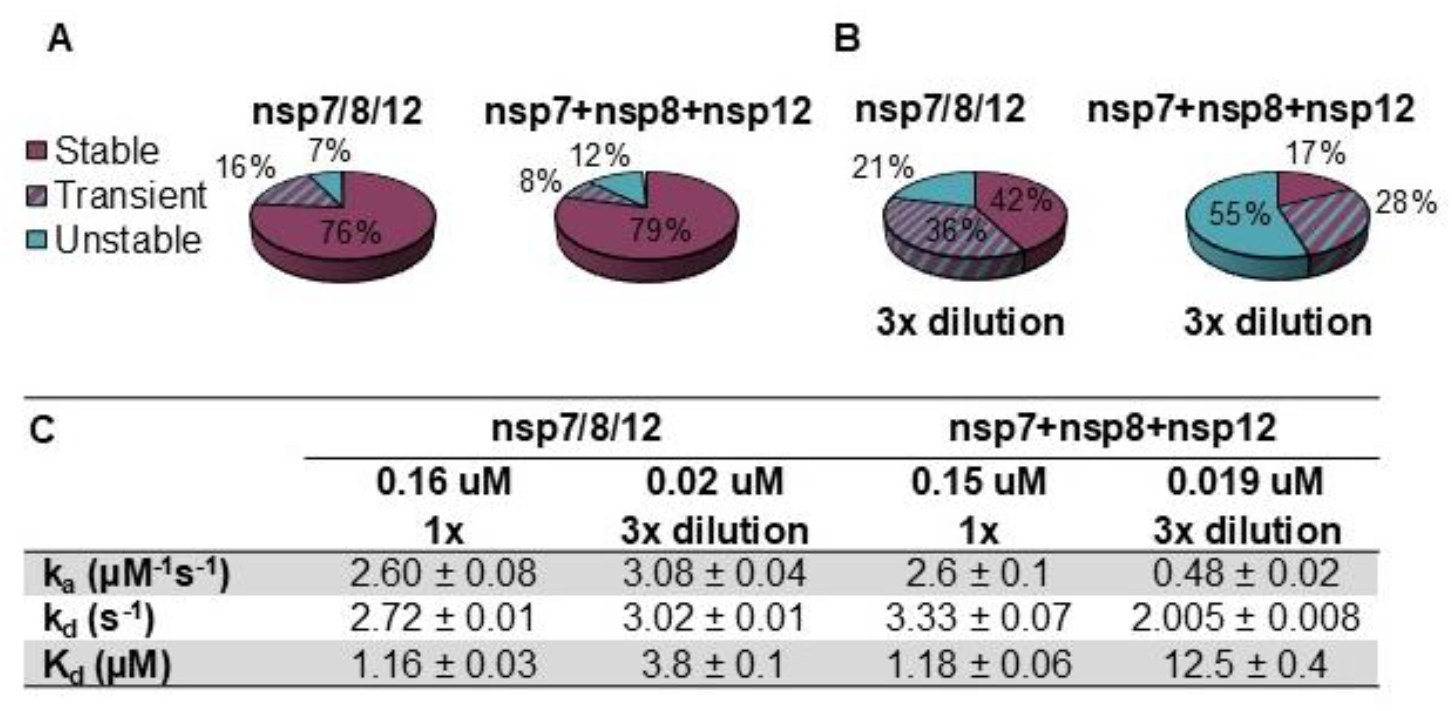
(**A**) Percent of protein binding (PIFE) to RNA breakdown of co-expressed and reconstituted (2:2:1 nsp7:nsp8:nsp12) protein. (**B**) Percent of protein binding to RNA when protein concentration was decreased by 3-fold and (**C**) corresponding kinetics for experiments in **A** and **B**.

The RdRp complex of the co-expressed components exhibited higher levels of RNA synthesis than the reconstituted complexes at 5:5:1 and 10:10:1 stoichiometry of the individual components (**Figure 3B**). To investigate whether a short RNA primer may exacerbate this difference, we conducted the RNA synthesis assay with a short, 4-nucleotide, RNA primer and a 20-nucleotide RNA template (**Figure S1**). Indeed, the reconstituted complexes failed to show measurable RNA synthesis under these conditions. This suggested that there may be a fundamental difference between protein production methods and the RdRp complexes of co-expressed components possess superior RNA synthesis activity. Altogether, these findings corroborate a relationship between stable binding observed and RdRp activity recorded.

### SM-assay distinguishes between inhibitor classes and reveals drug effects on polymerase complex binding

To validate the ability of the PIFE assay to differentiate between nucleoside analog polymerase inhibitors (NAPIs) and non-nucleosidic inhibitors (NNIs), binding of both separately expressed and co-expressed nsp7, nsp8, and nsp12 were assessed in the presence of suramin and remdesivir, respectively (**Figure 5**). NAPIs function as nucleotides but prevent further base incorporation thereby terminating RNA replication (**Figure 5A**, remdesivir triphosphate). On the other hand, NNIs hinder replication by blocking the polymerase docking site (**Figure 5A**, suramin). When either the separately or co-expressed nsps are combined with high remdesivir concentrations, the percentage of binding remained intact, and the ratio of stable to unstable binding did not change significantly. This suggests that remdesivir does not hinder RdRp binding to the RNA duplex, as expected. However, remdesivir does inhibit RNA synthesis (**Figure 3B**).

**Figure 5.**
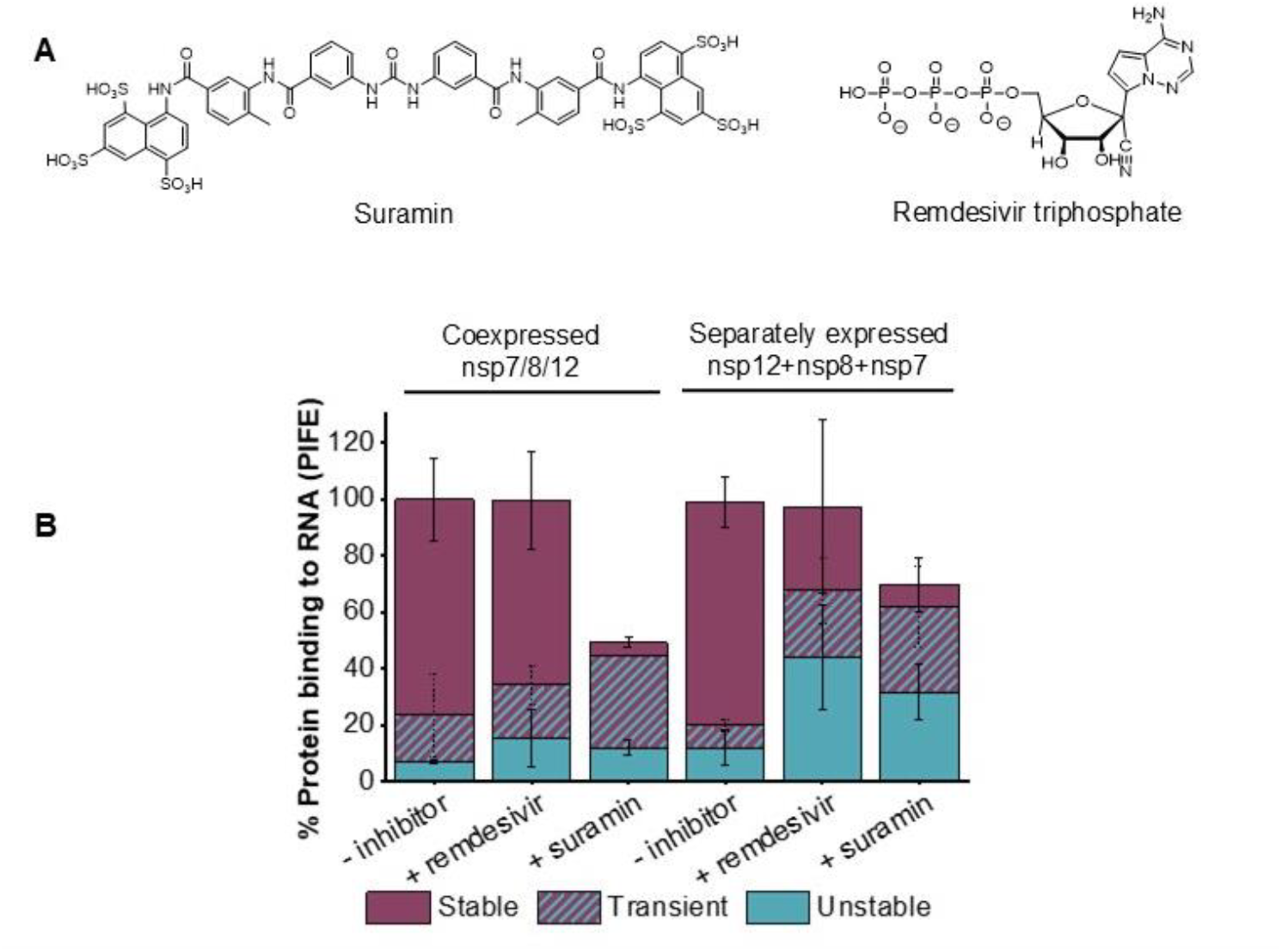
(**A**) Structures of suramin and remdesivir triphosphate. (**B**) Percent of molecules that showed protein binding to RNA (PIFE) broken into unstable and stable bindings when nsp7/8/12 was incubated in the absence of inhibitors, with 10 μM remdesivir, and with 5 μM suramin; showing the ability of the assay to differentiate between different inhibitor classes.

Suramin is well-known for its capability to competitively inhibit protein binding to nucleic acid substrates.(20) When the co-expressed nsp7/8/12 complex was exposed to suramin (5 µM), the proportion of molecules exhibiting PIFE was reduced from 100% to 44% (**Figure 5B**). Interestingly, not only was there a decrease in protein binding in the presence of suramin, but there was also a difference in the type of binding observed. With suramin addition, of the molecules that exhibit PIFE the amount of stable binding dropped significantly from 77% to 3%. This is likely because although suramin is in the RdRp binding pocket inhibiting productive binding, the protein can still bind to the RNA in a non-productive manner. To understand this further, the order of suramin addition was explored.

Pre-incubating the protein and RNA followed by suramin addition resulted in 64% stable binding, equivalent to the level observed in the absence of suramin (**Figure 6**). This suggests that the protein is not displaced by suramin, and further that the protein binding is tight, i.e., when the protein binds to RNA productively, it will not dissociate and be trapped by the suramin in solution. Conversely, pre-incubating the protein and suramin led to a notable increase in unstable bindings and a decrease in stable events. This illustrates that suramin had sufficient time to bind to the RdRp complex, thereby preventing the majority of RdRps from productively binding to RNA.

**Figure 6.**
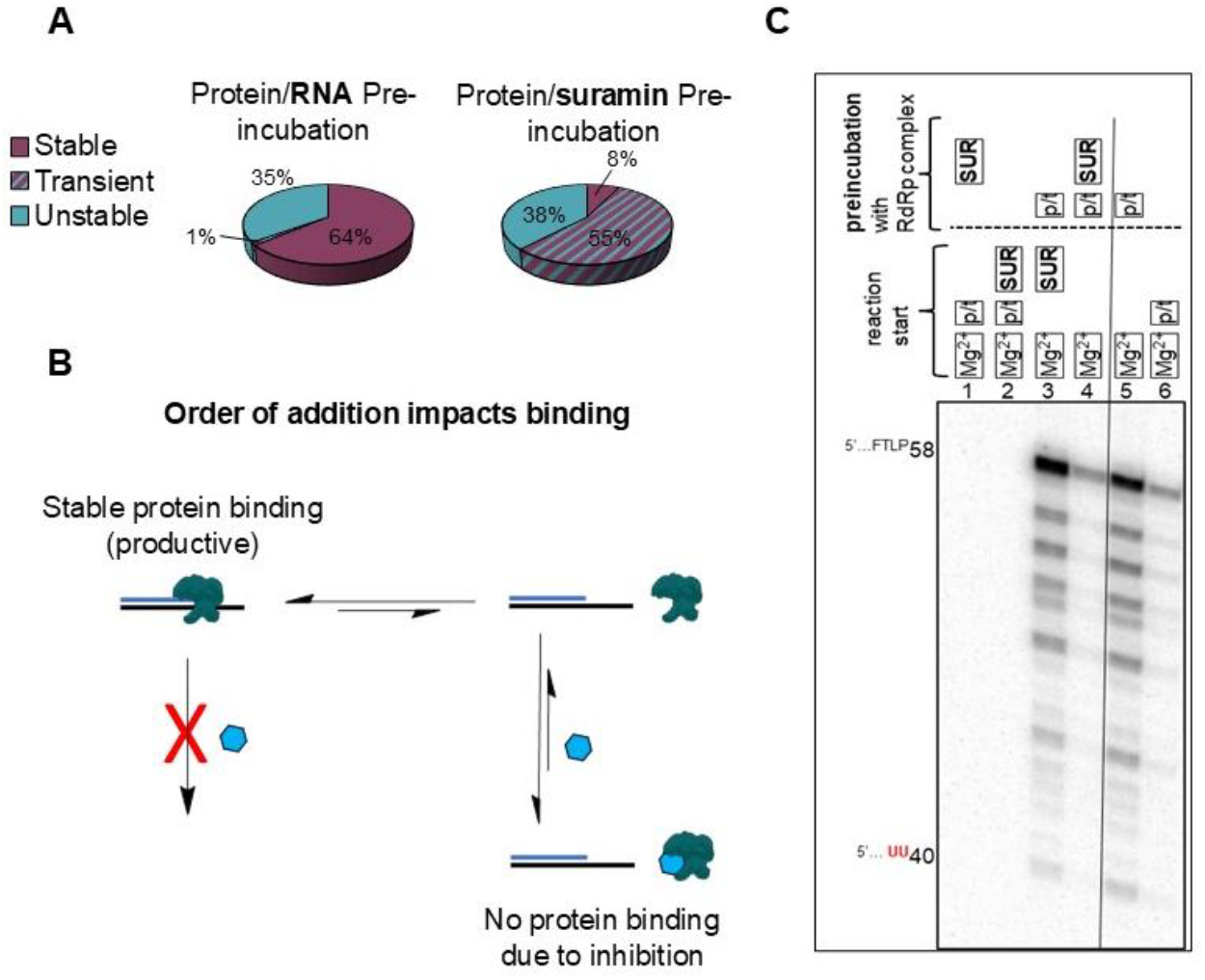
(**A**) % PIFE broken down by binding types when the RdRp (expressed as a polyprotein) is either pre-incubated with RNA or suramin to show how order of addition impacts binding. (**B**) Cartoon representing the dynamics of protein-RNA interactions in the presence of suramin with different order of additions, which is supported by (**C**) The effect of the order of addition on the inhibition of RNA synthesis by suramin. RNA synthesis reactions were conducted and the experimental readout was annotated as in Figure 3, except that order of addition of the reaction components during the preincubation versus the reaction start was varied. All reaction mixtures contained [α-^32^P]-UTP, ATP, CTP, and GTP mix and 10 µM suramin. The composition of the reaction components during the preincubation and the reaction start steps are illustrated in the panel above the gel. SUR, suramin. p/t, RNA primer/template, Mg^2+^, MgCl_2_. Numbers on top of the gel enumerate the gel lanes from left to right.

These dynamics outlined in **Figure 6B** are consistent with RNA synthesis assays (**Figure 6C**). Preincubating the RdRp complex with RNA primer/template resulted in higher levels of RNA synthesis activity as compared with omitting it from the preincubation step (**Figure 6C**, lanes 5 versus 6) illustrating that formation of the RdRp:RNA complex during the preincubation step promotes RNA synthesis. In line with this observation, supplementing the reactions with suramin and following the same order of addition as in *lane 6* completely inhibited RNA synthesis regardless of whether or not suramin was added or not during the preincubation step (**Figure** 6C, *lanes 1 and 2*). Allowing the RdRp:RNA complex to form during the preincubation step made it resistant to inhibition by suramin (**Figure 6C**, *lanes 5* versus *3*). However, including suramin during the preincubation step alongside RdRp complex and RNA dramatically reduced the RNA synthesis (**Figure 6C**, *lanes 5* versus *4*). This is consistent with suramin competing with RNA duplex for binding to RdRp complex. On the other hand, the visible remaining RNA synthesis activity in *lane 4* suggests that once the RdRp:RNA complex is formed it is stable enough to withstand the suramin challenge during the preincubation step.

## Conclusions

Single molecule PIFE studies revealed subpopulations of unstable and stable bindings of the SARS-CoV-2 RdRp with RNA duplexes. The opportunity to control order of addition enabled assessing the impact that each of the nsp7, 8 and 12 have in the overall assembly. Both nsp8 and nsp12 are responsible for efficient binding of the complex to its RNA duplex, while nsp7 has no impact on binding, although it is required for synthesis. While co-expressed protein showed stable binding, reconstituted mixtures displayed a range of unstable, transient and stable binding, indicating the molecular heterogeneity of these mixtures. The SM-PIFE assay, reporting on RdRp binding, aligns with the RNA synthesis gel-based assays - where enhanced binding recorded in SMPIFE studies is paralleled by enhanced RNA synthesys – while also providing detail on the dynamics between the nsps and RNA.

The ability to visualize and decipher the binding dynamics of key nsps opens new avenues for postulating dynamic/stoichiometric models for understanding replication mechanisms of emerging viruses. In turn the ability to screen newly developed inhibitors by controlling order of addition facilitates elucidating their mechanisms of action, a most valuable contribution to the field of antiviral drug discovery. As the COVID-19 pandemic continues to impact global health, these findings may aid in the development of more effective therapeutic interventions to combat viral infections.

## Data availability

All processed data for this study are included in this article and its Supporting Information. Movies and single molecule traces can be provided upon request.

## Conflict of interest statement

None declared.

## Acknowledgements

G.C. and M.G. thank NIH Rapidly emerging antiviral drug development initiative - AVIDD center (READDI-AC) Award #: U19AI171292. G.C. is thankful to the National Science and Engineering Research Council of Canada (NSERC), the Canadian Foundation for Innovation (CFI), the Fonds de Recherche du Québec – Nature et Technologies (FRQNT), and the Canada Institute for Health Research (CIHR) for funding. G.C. is also thankful to the Canada Research Chairs Program. M.G is also thankful for funding from SPP-ARC. T.C.L. acknowledges NSERC Postdoctoral Fellowship program (PDF) for funding. H.A.F.D. is thankful to NSERC CREATE PROMOTE for a postgraduate scholarship. C.J.G. is supported by a grant from the CIHR under the funding reference number 181545. The authors would like to thank Emma Woolner for excellent technical assistance and Dr Jack Moore at the Alberta Proteomics and Mass Spectrometry facility for mass spectrometry analysis.

## Notes

### Competing Interest Statement

The authors have declared no competing interest.

