## Supporting Information for "Single-molecule assay reveals the impact of composition, RNA duplex, and inhibitors on the binding dynamics of SARS-CoV-2 polymerase complex"

### 1. MATERIALS AND METHODS

#### 1.1 Materials.

**Slide Preparation.** Acetone (HPLC grade), Premium Cover Glass slides (25x25 mm), sulfuric acid, and sodium hydroxide were purchased from Fisher. Dimethyl sulfoxide (anhydrous, ≥99.9), Wheaton Coplin Staining Jars were purchased from SigmaAldrich. Hydrogen peroxide (30% v/v) was purchased from ACP Chemicals. Silicon molds, polycarbonate imaging chambers (Hybriwell™), and ports (press fit tubing connectors) were purchased from Grace Bio-Labs, Inc. Double sided tape to affix ports was purchased from Scor-pal. Poly(ethylene glycol) polymers mPEG-Silane (MW 5000) and Biotin-PEG-Silane (MW 5000) were purchased from Laysan Bio Inc. Polymers were dissolved in anhydrous dimethyl sulfoxide (DMSO) and a 25% (w/w) PEG-Silane and 0.25% (w/w) Bio-PEG-Sil solution was made.

**Nucleic Acids.** RNA constructs were purchased from Integrated DNA Technologies. Template and primer strands were annealed by incubating in the thermal cycler at 95ºC for 2 min followed by a gradual cooling step of 2ºC/min to 25ºC. The RNA construct with the longer 25 nt overhang (construct 1, below) consisted of template strand Cy3-tagged 55mer (/5Cy3N/GCA-CUU-AGA-UAU-GAC-UCG-UUC-UGC-AGG-CCA-GUU-AAU-AAC-GUC-UAA-GAC-ACA-GAU-C/3) and primer strand biotinylated 30mer (/5Bio/GAU-CUG-UGU-CUU-AGA-CGU-UAU-UAA-CUG-GCC/3). The RNA construct with the shorter 20 nt overhang (construct 2, below) consisted of template strand Cy3-tagged 58mer (/5Cy3N/CCA-CAC-AAC-ACC-UAC-GGC-AAU-GGA-GCG-CUG-GCA-GCG-GUU-AAC-GUC-UAA-GAC-ACA-GAU-C/3) and primer strand biotinylated 38mer (/5Bio/GAU-CUG-UGU-CUU-AGA-CGU-UAA-CCG-CUG-CCA-GCG-CUC-CA/3). The hybridized constructs are represented below.

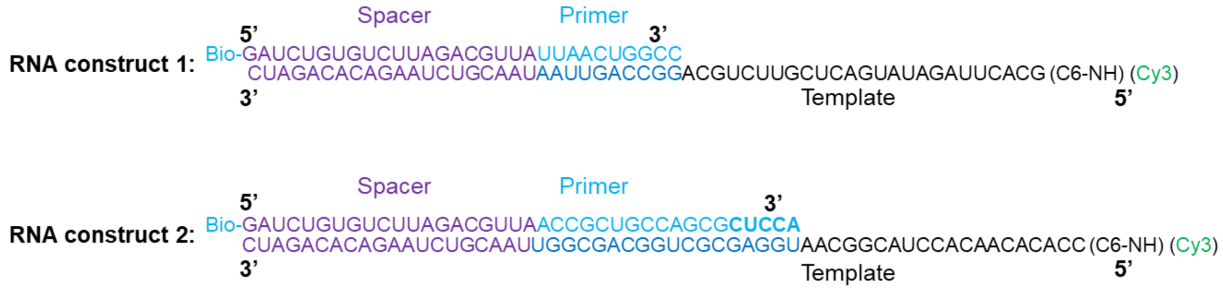

**Proteins.** All proteins were provided by Prof. Matthias Gotte at the University of Alberta.

**Buffers and reagents.** $MgCl_2$ 1.0 M, HEPES 1.0 M buffer solution pH 8.0, and Molecular Biology Grade HyClone were acquired from ThermoFisher Scientific (South Logan, UT). Unlabeled streptavidin (S888) and nucleoside triphosphate (CTP, ATP, GTP and UTP) 100 mM solutions were purchased from Thermofisher Scientific (Invitrogen). D-(+)-glucose, glucose oxidase (from Aspergillus niger, type VII) and biotin (HPLC grade lyophilized powder) were purchased from Sigma-Aldrich. Suramin was purchased from EMD Millipore. Remdesivir (active triphosphate metabolite GS-443902 trisodium) was purchased from MedChemExpress. Glycerol monothioglycolate (GMTG), 80%, was purchased from Evans Chemetics.

### 2. IMAGING AND DATA ANALYSIS.

An initial movie was acquired for 240 frames (50 ms exposure) in the presence of the anti-fading solution without protein. This is to acquire a time zero to localize the fluorophores and obtain a baseline fluorescence intensity before protein interaction with the RNA. Then the laser was turned off, a new anti-fading solution containing the protein of interest was injected, and a second movie was taken for 20,000 frames (50 ms exposure) to allow substantial time for the molecules to photobleach. The two movies were combined and aligned with ImageJ to correct drifting. Typically, ~400-800 single molecules over one field of view (~82 x 82 µm$^2$) were observed. Fluorescence intensity vs time trajectories of individual molecules were extracted from the acquired videos using an algorithm written in MATLAB (MathWorks) defining a region of interest around the centers of emission for the single molecules recorded. These molecules were picked by an intensity threshold. Intensity vs time trajectories were analyzed to determine the bound and unbound times for each molecule, where a sudden substantial intensity increase was determined to be a binding event and unbinding occurred once the intensity returned to the baseline. This data allowed plotting frequency histograms for the on time, off time, and photobleaching time. Fitting these histograms with a monoexponential decay function allowed determination of the average time for each event. The association constant ($k_a$) was calculated using the average unbound time ($\tau_{unbound}$) and RdRp concentration, as seen in **Equation 1**. The concentration used in the calculation varied based on the components present in the solution. When nsp12 was present, its concentration was employed in the calculation (0.15 μM unless otherwise stated), as it is the primary component of interest. In instances where nsp12 was absent and nsp8 was in solution, the concentration of nsp8 was utilized as it showed the most significant binding of the two accessory components. In cases where only nsp7 was present, its concentration was used for the calculation. The dissociation constant ($k_d$) was calculated using the average bound time ($\tau_{bound}$), as seen in **Equation 2**. The equilibrium dissociation constant ($K_d$) was calculated as seen in **Equation 3**. The percent of molecules with PIFE were manually counted. Stable binding is defined as binding events that are at least 10 times longer than the average unstable bound time. Unstable binding events are those that show protein unbind from the RNA. Transient binding are molecules that initially showed unstable events before achieving stable binding. Non-binding refers to molecules that showed no change in fluorescence intensity throughout the experiment. The number of molecules that were analyzed for each experiment (n) are also listed. Values are listed for three trials. Large discrepancies between trials result from protein degradation over time or differences in protein batches. Despite these variations, the main conclusions remain consistent.

### 3. TRIPLICATE KINETIC PARAMETERS AND PERCENT PIFE.

**Table S1**. Percent PIFE of the separately expressed nsp7, nsp8, and nsp12 components in a ratio of 1:2:1 immediately after protein addition and then after 30 minutes in the dark.

| Time | Binding type | Trial 1 | Trial 2 | Trial 3 | Average |
|---|---|---|---|---|---|
| **Initial** | Overall % PIFE | 99% | 97% | 100% | 99.8 ± 1.4% |
| | % Stable | 49% | 40% | 62% | 50 ± 11% |
| | % Unstable | 50% | 57% | 38% | 48 ± 10% |
| | % Non-binding | 1% | 3% | 0% | 1 ± 1% |
| | n | 250 | 289 | 237 | - |
| **After 30 minutes** | Overall % PIFE | 100% | 100% | 100% | 100 ± 0 % |
| | % Stable | 93% | 84% | 88% | 89 ± 5% |
| | % Unstable | 7% | 15% | 12% | 11 ± 4% |
| | % Non-binding | 0% | 0.5% | 0% | 0.2 ± 0.3% |
| | n | 174 | 177 | 136 | - |

**Table S2**. Percent PIFE and kinetic values for experiments with RNA construct 2 with coexpressed (nsp7/8/12) and separately expressed nsps.

| nsp component | Binding type | Trial 1 | Trial 2 | Trial 3 | Average |
|---|---|---|---|---|---|
| **nsp7/8/12** | % Overall PIFE | 100% | 99% | 100% | 100 ± 0% |
| | % Stable | 61% | 78% | 90% | 76 ± 15% |
| | % Transient | 32% | 14% | 3% | 16 ± 15% |
| | % Unstable | 7% | 8% | 7% | 7 ± 1% |
| | % Non-binding | 0.0% | 0.1% | 0.5% | 0.2 ± 0.3% |
| | n | 246 | 811 | 183 | - |
| | Unstable $\tau_{bound}$ (s) | 0.407 ± 0.001 | 0.727 ± 0.006 | 0.231 ± 0.002 | 0.455 ± 0.002 |
| | $\tau_{photobleaching}$ (s) | 117 ± 4 | 93 ± 10 | 64 ± 7 | 91 ± 4 |
| | $\tau_{unbound}$ (s) | 3.6 ± 0.1 | 1.8 ± 0.1 | 2.4 ± 0.1 | 2.60 ± 0.06 |
| | $k_a$ ($\mu M^{-1}s^{-1}$) | 1.74 ± 0.05 | 3.5 ± 0.2 | 2.6 ± 0.1 | 2.60 ± 0.08 |
| | $k_d$ ($s^{-1}$) | 2.457 ± 0.006 | 1.38 ± 0.01 | 4.33 ± 0.04 | 2.72 ± 0.01 |
| | $K_d$ (μM) | 1.42 ± 0.04 | 0.40 ± 0.02 | 1.66 ± 0.07 | 1.16 ± 0.03 |
| **nsp7** | % Overall PIFE | 4% | 17% | 5% | 9 ± 7% |
| | % Stable | 3% | 2% | 0% | 2 ± 1% |
| | % Transient | 1% | 4% | 0% | 2 ± 2% |
| | % Unstable | 2% | 10% | 5% | 6 ± 4% |
| | % Non-binding | 96% | 84% | 95% | 91 ± 7% |
| | n | 646 | 266 | 286 | - |
| | Unstable $\tau_{bound}$ (s) | 0.53 ± 0.05 (n=11) | 0.65 ± 0.09 (n=15) | 0.42 ± 0.02 (n=27) | 0.53 ± 0.04 |
| | $\tau_{photobleaching}$ (s) | 330 ± 98 | 65 ± 23 (n=28) | 326 ± 2 | 240 ± 34 |
| | $\tau_{unbound}$ (s) | 24.4 ± 3 | Too few data | 18.6 ± 0.9 | 22 ± 2 |
| | $k_a$ ($\mu M^{-1}s^{-1}$) | 0.055 ± 0.007 | N/A | 0.0717 ± 0.003 | 0.063 ± 0.003 |
| | $k_d$ ($s^{-1}$) | 1.9 ± 0.2 | 1.5 ± 0.2 | 2.4 ± 0.1 | 1.9 ± 0.1 |
| | $K_d$ (μM) | 35 ± 5 | N/A | 33 ± 2 | 34 ± 2 |
| **nsp8** | % Overall PIFE | 55% | 93% | 58% | 69 ± 21% |
| | % Stable | 4% | 9% | 0% | 4 ± 4% |
| | % Transient | 8% | 31% | 1% | 13 ± 16% |
| | % Unstable | 43% | 53% | 58% | 51 ± 8% |
| | % Non-binding | 45% | 7% | 42% | 31 ± 21% |
| | n | 263 | 302 | 146 | - |
| | Unstable $\tau_{bound}$ (s) | 0.290 ± 0.002 | 0.503 ± 0.004 | 0.512 ± 0.005 | 0.435 ± 0.002 |
| | $\tau_{photobleaching}$ (s) | 205 ± 58 | 559 ± 23 | 209 ± 10 | 324 ± 21 |
| | $\tau_{unbound}$ (s) | 22 ± 1 | 30 ± 2 | 12.0 ± 0.1 | 21.3 ± 0.7 |
| | $k_a$ ($\mu M^{-1}s^{-1}$) | 0.061 ± 0.003 | 0.044 ± 0.003 | 0.111 ± 0.001 | 0.072 ± 0.001 |
| | $k_d$ ($s^{-1}$) | 3.45 ± 0.02 | 1.99 ± 0.02 | 1.95 ± 2 | 2.46 ± 0.01 |
| | $K_d$ (μM) | 57 ± 3 | 45 ± 3 | 17.6 ± 0.2 | 40 ± 1 |
| **nsp12** | % Overall PIFE | 63% | 96% | 99% | 86 ± 20% |
| | % Stable | 13% | 25% | 32% | 23 ± 10% |

| | | | | | |
|---|---|---|---|---|---|
| | % Transient | 19% | 47% | 45% | 37 ± 16% |
| | % Unstable | 33% | 25% | 22% | 26 ± 5% |
| | % Non-binding | 37% | 4% | 1% | 14 ± 20% |
| | n | 237 | 441 | 273 | - |
| | Unstable $\tau_{bound}$ (s) | 0.424 ± 0.004 | 0.434 ± 0.009 | 0.547 ± 0.004 | 0.468 ± 0.004 |
| | $\tau_{photobleaching}$ (s) | 86 ± 4 | 166 ± 23 | 178 ± 9 | 143 ± 8 |
| | $\tau_{unbound}$ (s) | 5.3 ± 0.1 | 7.0 ± 0.6 | 7.6 ± 0.2 | 6.6 ± 0.2 |
| | $k_a$ ($\mu M^{-1}s^{-1}$) | 1.26 ± 0.02 | 0.95 ± 0.08 | 0.87 ± 0.02 | 1.03 ± 0.03 |
| | $k_d$ ($s^{-1}$) | 2.36 ± 0.02 | 2.30 ± 0.05 | 1.83 ± 0.01 | 2.16 ± 0.02 |
| | $K_d$ (μM) | 1.88 ± 0.04 | 2.4 ± 0.2 | 2.08 ± 0.06 | 2.13 ± 0.07 |
| **nsp7+nsp8** | % Overall PIFE | 93% | 64% | 94% | 84 ± 17% |
| | % Stable | 3% | 11% | 0% | 5 ± 6% |
| | % Transient | 19% | 14% | 8% | 14 ± 5% |
| | % Unstable | 72% | 39% | 86% | 65 ± 24% |
| | % Non-binding | 7% | 36% | 6% | 16 ± 17% |
| | n | 480 | 377 | 245 | - |
| | Unstable $\tau_{bound}$ (s) | 0.440 ± 0.003 | 0.500 ± 0.007 | 0.240 ± 0.002 | 0.393 ± 0.003 |
| | $\tau_{photobleaching}$ (s) | 575 ± 104 | 391 ± 28 | 439 ± 109 | 468 ± 51 |
| | $\tau_{unbound}$ (s) | 15.1 ± 0.2 | 20.8 ± 0.3 | 17 ± 1 | 17.6 ± 0.4 |
| | $k_a$ ($\mu M^{-1}s^{-1}$) | 0.088 ± 0.001 | 0.064 ± 0.001 | 0.078 ± 0.005 | 0.077 ± 0.002 |
| | $k_d$ ($s^{-1}$) | 2.27 ± 0.02 | 2.00 ± 0.03 | 4.17 ± 0.04 | 2.81 ± 0.02 |
| | $K_d$ (μM) | 25.7 ± 0.4 | 31.2 ± 0.6 | 53 ± 3 | 37 ± 1 |
| **nsp7+nsp12** | % Overall PIFE | 70% | 95% | 98% | 88 ± 15% |
| | % Stable | 48% | 34% | 65% | 49 ± 16% |
| | % Transient | 26% | 27% | 14% | 22 ± 7% |
| | % Unstable | 22% | 35% | 19% | 25 ± 8% |
| | % Non-binding | 30% | 5% | 2% | 12 ± 15% |
| | n | 585 | 564 | 384 | - |
| | Unstable $\tau_{bound}$ (s) | 0.510 ± 0.005 | 0.474 ± 0.005 | 0.478 ± 0.007 | 0.487 ± 0.003 |
| | $\tau_{photobleaching}$ (s) | 55 ± 5 | 66 ± 2 | 71 ± 4 | 64 ± 2 |
| | $\tau_{unbound}$ (s) | 4.06 ± 0.08 | 6.4 ± 0.1 | 6.4 ± 0.1 | 5.62 ± 0.05 |
| | $k_a$ ($\mu M^{-1}s^{-1}$) | 1.64 ± 0.03 | 1.04 ± 0.02 | 1.04 ± 0.02 | 1.24 ± 0.01 |
| | $k_d$ ($s^{-1}$) | 1.96 ± 0.02 | 2.11 ± 0.02 | 2.09 ± 0.03 | 2.05 ± 0.01 |
| | $K_d$ (μM) | 1.19 ± 0.03 | 2.03 ± 0.04 | 2.01 ± 0.04 | 1.74 ± 0.02 |
| **nsp8+nsp12** | % Overall PIFE | 95% | 94% | 100% | 96 ± 3% |
| | % Stable | 93% | 90% | 86% | 90 ± 4% |
| | % Transient | 5% | 5% | 9% | 6 ± 2% |
| | % Unstable | 2% | 5% | 5% | 4 ± 2% |
| | % Non-binding | 0.3% | 1.5% | 0.3% | 0.7 ± 0.7% |
| | n | 359 | 196 | 309 | - |
| | Unstable $\tau_{bound}$ (s) | 0.55 ± 0.01 | 1.16 ± 0.06 | 0.48 ± 0.02 | 0.73 ± 0.02 |

| | | | | | |
|---|---|---|---|---|---|
| | $\tau_{photobleaching}$ (s) | 61 ± 4 | 68 ± 5 | 61 ± 5 | 63 ± 3 |
| | $T_{unbound}$ (s) | 1.21 ± 0.05 | 2.6 ± 0.2 | 3.7 ± 0.3 | 2.5 ± 0.1 |
| | $k_a$ ($\mu M^{-1}s^{-1}$) | 5.5 ± 0.2 | 2.6 ± 0.2 | 1.8 ± 0.1 | 3.3 ± 0.01 |
| | $k_d$ ($s^{-1}$) | 1.82 ± 0.03 | 0.86 ± 0.05 | 2.08 ± 0.09 | 1.59 ± 0.03 |
| | $K_d$ (μM) | 0.33 ± 0.02 | 0.34 ± 0.03 | 1.2 ± 0.1 | 0.61 ± 0.04 |

**Table S3.** Percent PIFE of the separately expressed nsp7, nsp8, and nsp12 components in a ratio of 1:2:1 with RNA construct 1, which has a hairpin structure in the ss-RNA template overhang.

| **nsp component** | **Binding type** | **Trial 1** | **Trial 2** | **Trial 3** | **Average** |
|---|---|---|---|---|---|
| **nsp7+nsp8+nsp12 (1:2:1)** | % Overall PIFE | 84% | 99% | 85% | 90 ± 8% |
| | % Stable | 32% | 77% | 15% | 41 ± 32% |
| | % Transient | 21% | 18% | 35% | 25 ± 9% |
| | % Unstable | 31% | 5% | 36% | 24 ± 17% |
| | % Non-binding | 16% | 1% | 15% | 11 ± 8% |
| | n | 196 | 395 | 241 | - |

**Table S4**. Percent PIFE and kinetic values for experiments with RNA construct 2 and varying ratios of nsp7, nsp8, and nsp12.

| nsp7:nsp8:nsp12 ratio | Binding type | Trial 1 | Trial 2 | Trial 3 | Average |
|---|---|---|---|---|---|
| **(1:2:1)** | % Overall PIFE | 98% | 98% | 98% | 98 ± 0 |
| | % Stable | 73% | 77% | 87% | 79 ± 7% |
| | % Transient | 11% | 7% | 4% | 7± 4% |
| | % Unstable | 14% | 14% | 8% | 12 ± 4% |
| | % Non-binding | 2% | 2% | 2% | 2 ± 0% |
| | n | 551 | 624 | 377 | - |
| | Unstable $\tau_{bound}$ (s) | 0.65 ± 0.01 | 0.38 ± 0.01 | 0.172 ± 0.006 | 0.401 ± 0.005 |
| | $\tau_{photobleaching}$ (s) | 58 ± 12 | 52 ± 8 | 39 ± 8 | 50 ± 5 |
| | $T_{unbound}$ (s) | 3.2 ± 0.1 | 2.7 ± 0.1 | 2.0 ± 0.2 | 2.63 ± 0.08 |
| | $k_a$ ($\mu M^{-1}s^{-1}$) | 2.08 ± 0.07 | 2.47 ± 0.09 | 3.3 ± 0.3 | 2.6 ± 0.1 |
| | $k_d$ ($s^{-1}$) | 1.54 ± 0.02 | 2.63 ± 0.07 | 5.8 ± 0.2 | 3.33 ± 0.07 |
| | $K_d$ (μM) | 0.74 ± 0.03 | 1.07 ± 0.05 | 1.7 ± 0.2 | 1.18 ± 0.06 |
| **(3:3:1)** | % Overall PIFE | 100% | 92% | 93% | 95 ± 5% |
| | % Stable | 49% | 55% | 45% | 50 ± 5% |
| | % Transient | 35% | 16% | 9% | 20 ± 13% |
| | % Unstable | 17% | 21% | 38% | 25 ± 11% |
| | % Non-binding | 0% | 8% | 7% | 5 ± 5% |
| | n | 260 | 248 | 128 | - |
| | Unstable $\tau_{bound}$ (s) | 0.743 ± 0.005 | 0.446 ± 0.004 | 0.311 ± 0.007 | 0.500 ± 0.003 |
| | $\tau_{photobleaching}$ (s) | 188 ± 11 | 175 ± 23 | 37 ± 5 | 133 ± 9 |
| | $T_{unbound}$ (s) | 1.04 ± 0.02 | 4.20 ± 0.06 | 4.4 ± 0.3 | 3.2 ± 0.1 |
| | $k_a$ ($\mu M^{-1}s^{-1}$) | 6.4 ± 0.1 | 1.59 ± 0.02 | 1.5 ± 0.1 | 3.17 ± 0.05 |
| | $k_d$ ($s^{-1}$) | 1.346 ± 0.009 | 2.24 ± 0.02 | 3.22 ± 0.07 | 2.27 ± 0.03 |
| | $K_d$ (μM) | 0.210 ± 0.004 | 1.41 ± 0.02 | 2.1 ± 0.2 | 1.25 ± 0.05 |
| **(5:5:1)** | % Overall PIFE | 99% | 98% | 100% | 99 ± 1% |
| | % Stable | 75% | 73% | 89% | 79 ± 9% |
| | % Transient | 10% | 9% | 5% | 8 ± 2% |
| | % Unstable | 14% | 17% | 5% | 12 ± 6% |
| | % Non-binding | 2% | 1% | 0% | 1 ± 1% |
| | n | 526 | 687 | 312 | - |
| | Unstable $\tau_{bound}$ (s) | 0.43 ± 0.01 | 0.489 ± 0.005 | 0.409 ± 0.003 | 0.443 ± 0.004 |
| | $\tau_{photobleaching}$ (s) | 56 ± 6 | 69 ± 5 | 82 ± 8 | 69 ± 4 |
| | $T_{unbound}$ (s) | $\tau_{off}$ (s) | 1.65 ± 0.06 | 2.95 ± 0.09 | 2.89 ± 0.05 |
| | $k_a$ ($\mu M^{-1}s^{-1}$) | 4.0 ± 0.1 | 2.260 ± 0.069 | 2.31 ± 0.04 | 2.87 ± 0.05 |
| | $k_d$ ($s^{-1}$) | 2.33 ± 0.05 | 2.05 ± 0.02 | 2.45 ± 0.02 | 2.27 ± 0.02 |
| | $K_d$ (μM) | 0.58 ± 0.03 | 0.91 ± 0.03 | 1.06 ± 0.02 | 0.85 ± 0.01 |

**4. INHIBITOR PIFE RESULTS.**

**Table S5**. Percent PIFE for experiments with RNA construct 2 with coexpressed and separately expressed nsps in the presence of remdesivir (10 μM) or suramin (5 μM).

| nsp component | Binding type | Trial 1 | Trial 2 | Trial 3 | Average |
|---|---|---|---|---|---|
| **nsp7/8/12 + remdesivir** | % Overall PIFE | 100% | 100% | 99% | 100 ± 1% |
| | % Stable | 67% | 82% | 48% | 65 ± 17% |
| | % Transient | 20% | 11% | 25% | 19 ± 7% |
| | % Unstable | 13% | 7% | 27% | 16 ± 10% |
| | % Non-binding | 0.2% | 0.0% | 1.0% | 0.4 ± 0.6% |
| | n | 574 | 271 | 191 | - |
| **nsp7/8/12 + suramin** | % Overall PIFE | 48% | 50% | 33% | 44 ± 9% |
| | % Stable | 6% | 3% | 0% | 3 ± 3% |
| | % Transient | 33% | 33% | 7% | 24 ± 15% |
| | % Unstable | 10% | 14% | 26% | 17 ± 8% |
| | % Non-binding | 52% | 50% | 67% | 56 ± 9% |
| | n | 397 | 208 | 274 | - |
| **nsp7+nsp8+nsp12 + remdesivir** | % Overall PIFE | 99% | 97% | 97% | 97 ± 1% |
| | % Stable | 65% | 17% | 7% | 30 ± 31% |
| | % Transient | 10% | 32% | 28% | 23 ± 12% |
| | % Unstable | 24% | 47% | 61% | 44 ± 19% |
| | % Non-binding | 1% | 4% | 4% | 3 ± 1% |
| | n | 434 | 597 | 426 | - |
| **nsp7+nsp8+nsp12 + suramin** | % Overall PIFE | 80% | 60% | 70% | 70 ± 10% |
| | % Stable | 15% | 1% | 2% | 6 ± 8% |
| | % Transient | 40% | 20% | 8% | 23 ± 16% |
| | % Unstable | 25% | 39% | 60% | 41± 18% |
| | % Non-binding | 20% | 40% | 30% | 30 ± 10% |
| | n | 426 | 726 | 450 | - |

### 5. GEL-BASED POLYMERIZATION ASSAYS

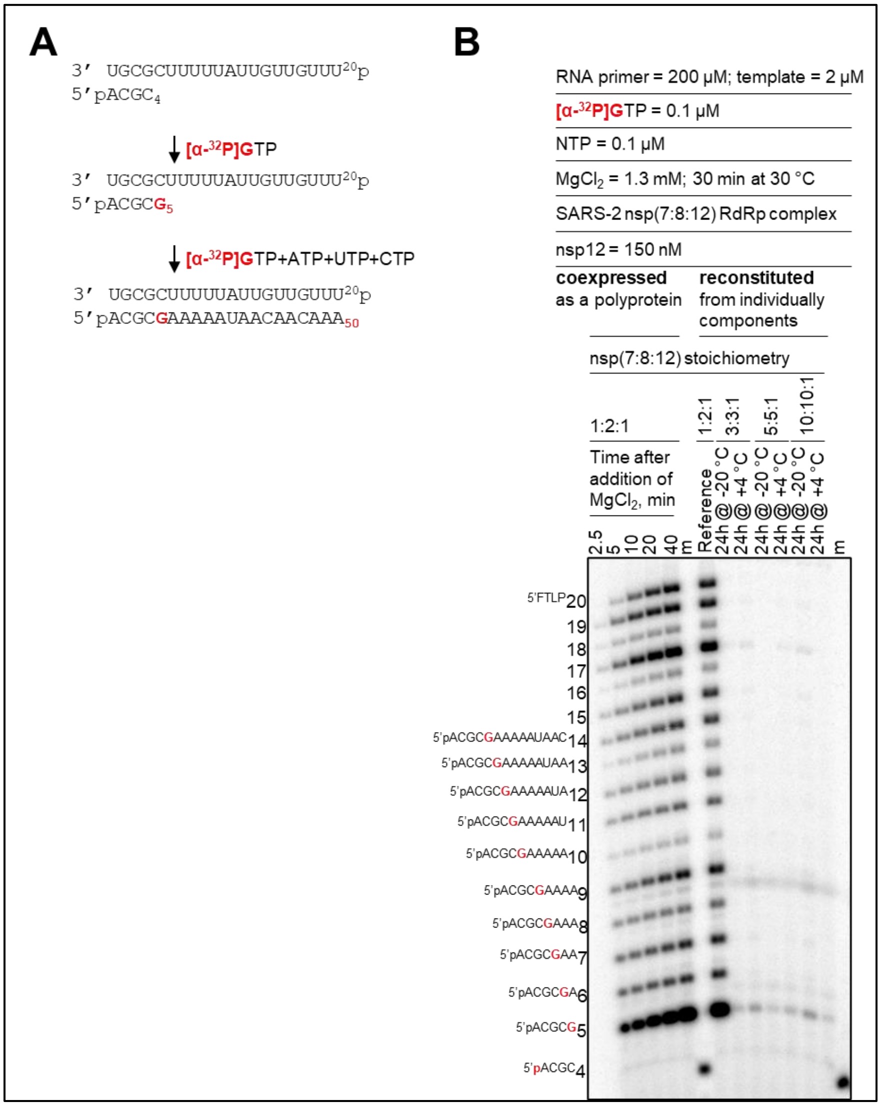

**Figure S1. Comparison of RNA synthesis activity on short RNA primer/RNA template catalyzed by SARS-CoV-2 RdRp complexes expressed as a viral polyprotein or reconstituted from individual components.**

(**A**) Schematic representation of the RNA synthesis assay. Model RNA template and a short RNA primer system used in the RNA synthesis assays. Numbers in superscript format indicate the nucleotide length of the template. Numbers in subscript format indicate the nucleotide length of the primer and the expected products. Downward arrows illustrate the addition of specific NTP mixes to the reaction mixtures, which allows the formation of primer extension products with defined length. *Bold red*, illustrates a radiolabeled GTP added to the reaction mixture, or incorporated radiolabeled GMP during primer extension, which effectively labels the reaction products and allows signal detection and quantification. (**B**) Denaturing PAGE migration pattern of the products of RNA synthesis reactions at indicated concentrations and along the primer/template shown in panel A. The formation of the reconstituted RdRp complexes was allowed to proceed for 24 hours at 4 °C or for 30 min on ice and then stored at -20 °C. RNA synthesis assay consisted of pre-incubating the indicated RdRp complexes with the RNA primer and RNA template in the presence of [α-$^{32}$P]-GTP, and the indicated NTP mix for 10 minutes at

30 °C. The RNA synthesis was initiated by addition of $MgCl_2$. Reactions mixtures were incubated for indicated time points or 40 minutes at 30 °C and then stopped with denaturing gel-loading buffer and subjected to denaturing polyacrylamide gel electrophoresis to resolve products of RNA synthesis. NTP, a mixture of ATP, CTP, and UTP. FTLP, full template-length product. m, 5'-radiolabeled 4-nucleotide marker. Sequences of letters shown to the left of the gel illustrate the 4-nucleotide marker or the nucleotide-extended primers in the presence of [α-$^{32}$P]-GTP and the NTP mix added to the reaction mixtures as per panel A. Numbers associated with the letter sequences illustrate the nucleotide size of the marker or the extended primers.
